# Synchronisation of yeast cell cycle through quorum sensing coupling

**DOI:** 10.1101/2020.04.05.026179

**Authors:** Giansimone Perrino, Diego di Bernardo

## Abstract

The cell cycle is present in all cells of all species and it is of fundamental importance in coordinating all the steps required for cell replication, including growth, DNA replication and cell division. Budding yeast is an unicellular organism characterised by a mother cell that buds to generate a daughter cell at each cell cycle. Each cell in a population buds at a different time. Despite its importance in biological applications, such as unravelling cell-cycle machinery mechanisms and production of valuable bioproducts, at present no yeast strain is capable of budding synchronously. To overcome this problem, we used control theory to propose a strategy to modify the yeast cell to endow it with the ability to synchronise its cell cycle across the population. Our strategy relies on a quorum sensing molecule diffusing freely in and out of the cell. The quorum sensing molecule is produced only during a specific phase of the cell cycle and couples the cell-cycle across the cell population. Here we model the proposed strategy with ordinary differential equations and numerically simulate it to demonstrate the feasibility of such an approach.

## 1. INTRODUCTION

Biological clocks give rise to rhythmic behaviour that are of fundamental importance in the coordination of many life processes (Winfree, 1967). One of the most prominent examples is the cell cycle machinery that generates rhythms that coordinate the vital process of cell replication by division, essential to guarantee the survival of species.

Cell cycle is a natural phenomenon occurring asynchronously in budding yeast *Saccharomyces cerevisiae* and whose regulation is highly conserved among the eukaryotes (Li et al., 2004). The cell cycle process consists of four sequential phases: G1, S (synthesis phase), G2, and M (mitosis phase). The cell grows in size during the G1 phase. Then, the G1 checkpoint mechanism triggers the start of the cell division process, only if a set of cellular and environmental conditions favourable to cell replication is satisfied. Upon G1 checkpoint release, the next three phases of cell cycle occur, ending with cell division at the end of the M phase, then the cell re-enters the G1 phase, thus completing the cycle. The overall process can be described by the state diagram illustrated in Fig. 1.

**Fig. 1.**
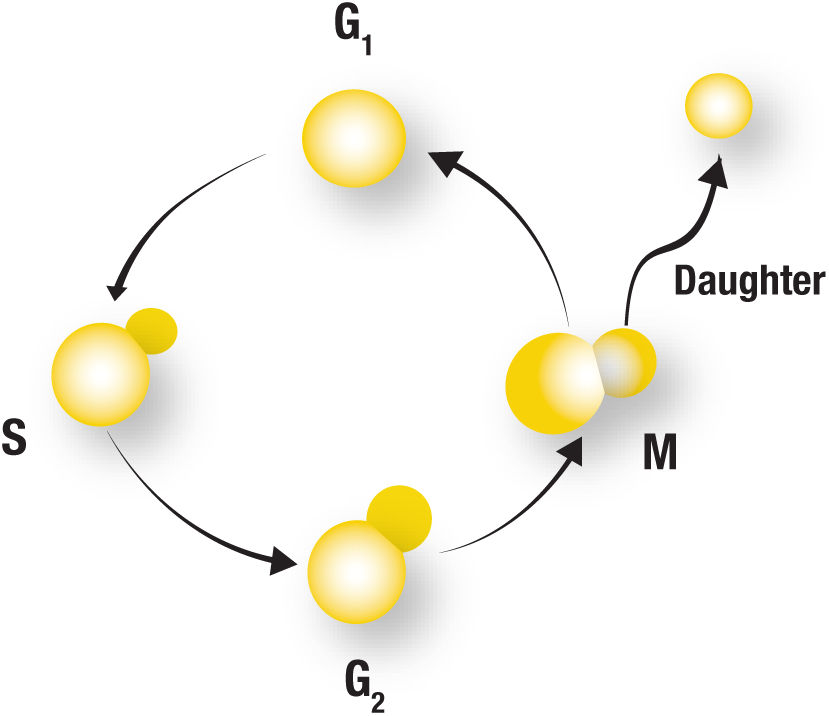
Diagram of cell cycle progression in yeast *Saccharomyces cerevisiae*.

Normally, cell division events occur asynchronously in the cell populations, meaning that cells do not bud at the same time. The reason behind this behaviour is that the cell-cycle clocks are not coupled across cells in the same population. From a biological point of view, a desynchronised collective behaviour could even be beneficial to increase cellular heterogeneity in the event of a catastrophe that can decimate the entire cell population (Acar et al., 2008; Beaumont et al., 2009).

However, there are situations in which it is desirable to have synchronised cell-cycle across the cell population, such as investigation of cell-cycle regulatory mechanisms (Davis et al., 2001), or production of valuable bio-products. On top of this, it is interesting to check whether feedback control theory can be applied also to such a complex system. Biologists developed different methods to force a cell population to divide synchronously (Davis et al., 2001). However, all of these methods do not dynamically synchronise the cell population, but just force each cell in the population to start from the same initial condition. Therefore, after a few generations the cell-cycle clocks are again desynchronised because of extrinsic and intrinsic biological noise (Di Talia et al., 2007).

Here, we propose a feedback control approach to synchronise the cell-cycle across the cell population. Specifically, we propose to synchronise the cell-cycle clocks in budding yeast by coupling them through a cell-to-cell communication system based on the elements of the quorum sensing machinery of bacterium *Vibrio fischeri* (Miller and Bassler, 2001). We investigated the feasibility to endow cells with controllers exploiting cell-to-cell communication to synchronise the cell cycle across the cell population, through theoretical and numerical analyses.

Numerical results demonstrate the feasibility of building a synthetic biomolecular control circuit to solve the cell cycle synchronisation problem in living yeast cells. The approach presented here constitutes an example of biological application in the area of *Cybergenetics*, an interdisciplinary field that works at the interface between *Control Theory* and *Synthetic Biology* for engineering synthetic controllers able to steer the dynamics of living systems (Khammash et al., 2019).

## 2. SYNCHRONISED CELL-CYCLE CLOCKS THROUGH QUORUM SENSING COUPLING

Our control objective is to synchronise the cell-cycle clocks across a population of yeast cells. Basically, this means that all the cells have to be in the same phase during the time evolution. This also implies that cell division events occur synchronously.

To couple the cell-cycle process across cells, we consider endowing yeast cells with a cell-to-cell communication system based on the quorum sensing machinery of bacterium *Vibrio fischeri* (Miller and Bassler, 2001). In its natural implementation, the quorum sensing system produces and releases in the extracellular environment a signalling small molecule, here defined as an inducer molecule, that increases in concentration as a function of cell density (Danino et al., 2010). The inducer molecule can freely diffuse in and out of the cell and once inside the cell can activate the production of downstream genes via specific inducer-sensitive promoters. We thus propose to incorporate this cell-to-cell signalling mechanism in the cell cycle machinery to couple this process across cells, as shown in Fig. 2b. In our design, the inducer molecule is produced during the G1 phase and released into the extracellular medium. The inducer molecule can then be imported in the cell, and activate the start of the cell cycle, but only if the cell is in the G_1_ phase.

**Fig. 2.**
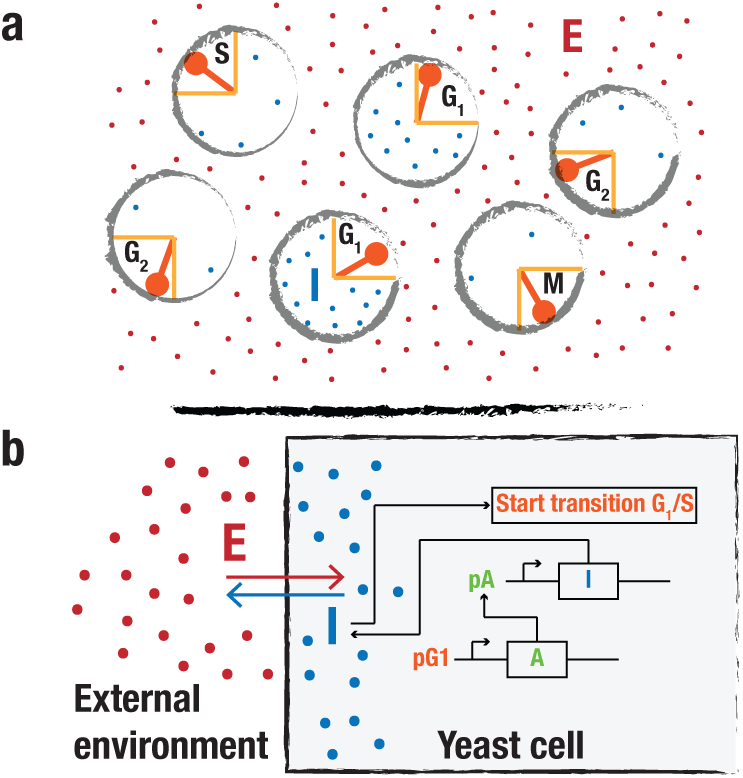
Quorum sensing coupling for synchronising cell-cycle across yeast cells. **a**, Schematics of the working principle of the quorum sensing coupling. **I** is the inducer molecule inside the cell, and it is produced only when a cell is in the G1 phase. The inducer molecule is free to diffuse in and out of the cell through a diffusion-like mechanism via the cell membrane. **E** is the inducer molecule in the extracellular environment. When the inducer molecule concentration inside the cell in the G1 phase reaches a threshold value, it triggers the transition to the synthesis (S) phase. **b**, Biological implementation of the synthetic biomolecular circuit coupling the cell cycle machinery in the cell population through the inducer molecule.

For the purpose of controlling the cell cycle progression by means of the quorum-sensing system, we need to modify the cell cycle machinery to enable its control by the inducer molecule. In budding yeast, three G1 cyclins (CLN1, CLN2, and CLN3) control the transition from the G1 phase to the S phase. However, expression of just one of these three cyclins is sufficient. Thus, we propose to delete the two endogenous G1 cyclins CLN1 and CLN3 from the genome, as done by Rahi et al. (2016), and drive *CLN2* gene expression from an inducible promoter sensitive to the quorum sensing inducer molecule **I**. At the same time, **I** is produced by an activator protein **A**, which is itself produced only in the G1 phase. Thus, in absence of inducer molecule, the cell-cycle is stuck in the G1 phase. However, in this phase, the inducer molecule is produced by the cell itself via the activator molecule and can freely diffuse into the extracellular environment (**E**), as illustrated in Fig. 2a. The addition of the quorum sensing mechanism thus effectively couples the cell cycle across cells and may induce synchronisation. In what follows we denote by *I*, and *E* the concentration of the signalling molecules inside the intracellular environment, and the extracellular environment, respectively.

## 3. MATHEMATICAL MODEL

We provide here a deterministic mathematical description of the considered biomolecular circuit presented in Section 2. Consider a population of *N* ∈ ℕ homogeneous yeast cells. First, we derive an ordinary differential equation describing the behaviour of the modified cell cycle machinery.

Mathematically, the cell cycle can be described as a dynamical system of the general form:

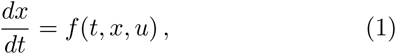

where *t* ∈ ℝ is time, *x* ∈ ℝ^*n*^ is the state vector, and *u* ∈ ℝ^*m*^ is the input vector of the possible external perturbations that affect the cell cycle machinery. If (1) has an exponentially stable limit cycle *γ* ⊂ ℝ^*n*^ with period *T*, then (1) is an oscillator that, according the phase reduction method (Winfree, 2001; Kuramoto, 1984), can be modelled as a dynamical phase oscillator

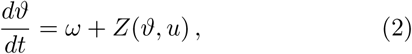

where *ϑ* is the phase of the oscillator on the unit circle 𝕊^1^, 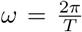 is the angular frequency, and *Z*: 𝕊^1^ × ℝ^*m*^ → ℝ is the 2*π*-periodic phase response curve (PRC) of the oscillatory system to *u* (Sacre and Sepulchre, 2014).

To model the phase evolution of the modified cell-cycle, we derive the following ordinary differential equation to describe the dynamics of cell-cycle phase of one cell (cell *i*):

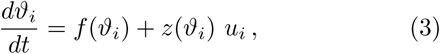

where *i* ∈ {1, …, *N*} denotes the *i*-th cell of the population, *ϑ*_*i*_ ∈ 𝕊^1^ is the 2*π*-periodic cell cycle phase, and *u*_*i*_ ∈ ℝ^+^ is the single bounded input acting on the cell cycle machinery via the internal inducer molecule *I*_*i*_. The phase-dependent switching function *f*: 𝕊^1^ → ℝ describes the fact that the cell-cycle stops in the G1 phase, if there is not enough inducer molecule (e.g. *u*_*i*_ = 0):

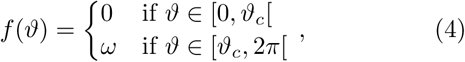

where *ϑ*_*c*_ ∈ 𝕊^1^ represents the end of the G_1_ phase and the beginning of the S phase. The phase response curve *z*: 𝕊^1^ → ℝ describes the linear response of the cell-cycle phase *ϑ*_*i*_ to the input *u*_*i*_:

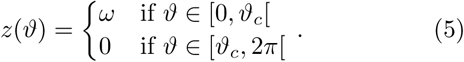

The input *u*_*i*_ is a Hill function of *I*_*i*_ with dissociation constant *K* and Hill coefficient *h*:

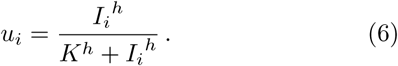

The dynamics of the activator *A* that couples the cell-cycle clock with the quorum sensing mechanism is described by the following ordinary differential equation:

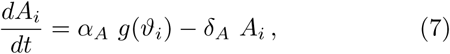

where *i* ∈ {1, …, *N*} is the *i*-th cell of the population, *α*_*A*_ is the production rate of species *A*_*i*_, *δ*_*A*_ is the degradation rate of species *A*_*i*_, and *g*: ℝ → ℝ is the 2*π*-periodic switching function that links the production of *A*_*i*_ to the cell cycle phase. Specifically, the species *A*_*i*_ is produced only when the cell cycle is in the G1 phase:

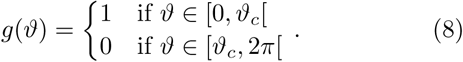

Next, we describe the dynamics of the inducer molecule in the intracellular and the extracellular environment. The molecules flow in and out of the extracellular environment through a diffusion-like phenomenon, hence the dynamics of *E* can be modelled using the partial differential equation:

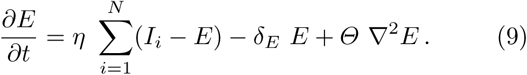

Equation 9 comprises three terms. The first one is the diffusion term between the intracellular and the extracellular environment, and it is characterised by the cell membrane diffusion rate *η*. The second term describes the degradation of the inducer molecule in the extracellular environment with degradation rate *δ*_*E*_. The third term represents the diffusion of the inducer molecules in the extracellular environment, and it is characterised by the external diffusion coefficient *Θ*. To solve the partial differential equation, we consider periodic boundary conditions in (9).

Notwithstanding the diffusion-like coupling requires a continuous spatial model to describe its own behaviour, the discrete spatial organisation of the cells inside the space allows the use of ordinary differential equations rather than partial differential equations (Ma and Yoshikawa, 2009). To discretise the partial differential equation (9), we derived a multi-compartment model by dividing the continuous spatial environment in as many compartments as the number of cells in the population, in this case *N*. The *i*-th cell is surrounded by the *i*-th compartment, and it is able to exchange the internal inducer molecules *I*_*i*_ only with its own compartment. Instead, the compartments are able to exchange the extracellular inducer molecules with their neighbours. We denote by *E*_*i*_ the discrete space concentration of the inducer molecules in the extracellular environment of the *i*-th compartment. The overall system can be seen as a network of *N* cells. Indeed, the compartments represent the nodes, whereas the exchange of the extracellular inducer molecules establishes the links among the nodes. The way in which the nodes are linked among them defines the topology of the network.

So, the discretisation of (9) relies on the assumptions made on the continuous spatial model and on the topology of the multi-compartment model. To this end, we consider a population of *N* cells in a one-dimensional space, where each cell exchanges information only with its left and right neighbour. Such assumption was similarly done by Danino et al. (2010), and it represents a simplification of the real quorum sensing system that well adapts to understand the collective behaviour of the population in a simplified manner. Hence, we replace (9) by a one-dimensional array of *N* ordinary differential equations representing individual cells coupled through the second-order discrete diffusion operator Δ*y*^−2^ (*E*_*i*−1_ + *E*_*i*+1_ − 2 *E*_*i*_), where *y* denotes the spatial coordinate of the one-dimensional space. Considering *D* = *Θ* Δ*y*^−2^ as the new external diffusion rate parameter, then the dynamics of the inducer molecules in the extracellular environment becomes:

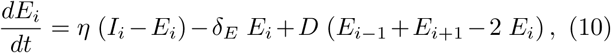

where *i* ∈ {1, …, *N*} is the *i*-th cell of the population. Also in this case we consider periodic boundary conditions at the limits of the array of *N* cells, that is *E*_0_ = *E*_*N*_ and *E*_*N*+1_ = *E*_1_. Under the latter assumption, the network of *N* cells is characterised by a ring topology.

Although the ring topology captures well a simplified quorum sensing communication, we also considered a more realistic scenario in which the compartments are free to communicate with all the other compartments, thus realising an all-to-all coupling among the cells. Assuming an all-to-all coupling topology, then (10) becomes:

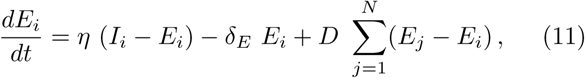

where *i* ∈ {1, …, *N*} is the *i*-th cell of the population. Finally, we modelled the dynamics of the intracellular inducer molecules *I* with the following ordinary differential equation:

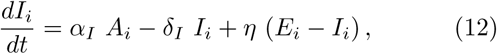

where *i* ∈ {1, …, *N*} is the *i*-th cell of the population, *α*_*I*_ is the production rate of *I*_*i*_, *δ*_*I*_ is the degradation rate of *I*_*i*_, and *η* is the cell membrane diffusion rate between the intracellular and the extracellular environment.

## 4. ANALYSIS OF COLLECTIVE BEHAVIOUR

Here, the synthetic quorum sensing coupling is simulated to assess its performance in synchronising the cell-cycle clocks. We provide the following definition:

### Definition 1. (Asymptotic phase synchronisation).

A population of *N* cells, modelled by (3), (7), (10), and (12); achieves asymptotic phase synchronisation if all the phases *ϑ*_*i*_(*t*) in (3) converge to the same value for *t* → ∞.

The same definition holds in the case of an all-to-all coupling, that is considering (11) in lieu of (10).

All numerical simulations were performed by considering a fixed population of *N* identical cells. The initial phases *ϑ*_*i*_(0) of individual cells were uniformly spaced in the interval [0, 2*π*[. The initial conditions of concentrations *I*_*i*_(0), *E*_*i*_(0), and *A*_*i*_(0) were all set equal to zero. Unless otherwise specified, numerical simulations were run with nominal parameters as reported in Table 1. The nominal parameters were chosen heuristically although they were coherent with the possible biological implementation of the overall system. Note that the concentrations are dimensionless. The system composed of the set of ordinary differential equations was solved using the MATLAB ode23t solver.

**Table 1.** Nominal parameter values used for the numerical simulations.

| Parameter | Value | Units |
| --- | --- | --- |
| $\omega$ | $0.025 \pi$ | $\text{rad} \cdot \text{min}^{-1}$ |
| $\vartheta_c$ | $0.5 \pi$ | rad |
| $K$ | 0.5 | |
| $h$ | 8 | |
| $\alpha_A$ | 20 | $\text{min}^{-1}$ |
| $\delta_A$ | 1 | $\text{min}^{-1}$ |
| $\alpha_I$ | 0.05 | $\text{min}^{-1}$ |
| $\delta_I$ | 1 | $\text{min}^{-1}$ |
| $\delta_E$ | 1 | $\text{min}^{-1}$ |
| $\eta$ | 2 | $\text{min}^{-1}$ |
| $D$ | 160 | $\text{min}^{-1}$ |

### 4.1 Numerical validation

We first ran a series of numerical simulations considering a small population of *N* = 10 cells with both ring and all-to-all topology. Although the number of cells is not biologically realistic, we decided to validate the proposed system first on networks of small dimensions, and then on networks composed by a larger number of nodes.

Numerical simulations show that asymptotic phase synchronisation of cell cycle clocks can be achieved with both topologies even when starting from a totally desynchronised population. Illustrative numerical simulations are reported in Fig. 3a (ring topology) and Fig. 3b (all-to-all topology). It can be appreciated that after a transient of approximately 25 hrs, the solutions of *ϑ*_*i*_, *I*_*i*_, and *E*_*i*_ reach a consensus value and become periodic. Also, the all-to-all coupling yields a faster transient achieving asymptotic phase synchronisation earlier than the ring coupling. Definitely, both topologies guarantee asymptotic phase synchronisation in at most 25 hrs, which is reasonable from an experimental point of view.

**Fig. 3.**
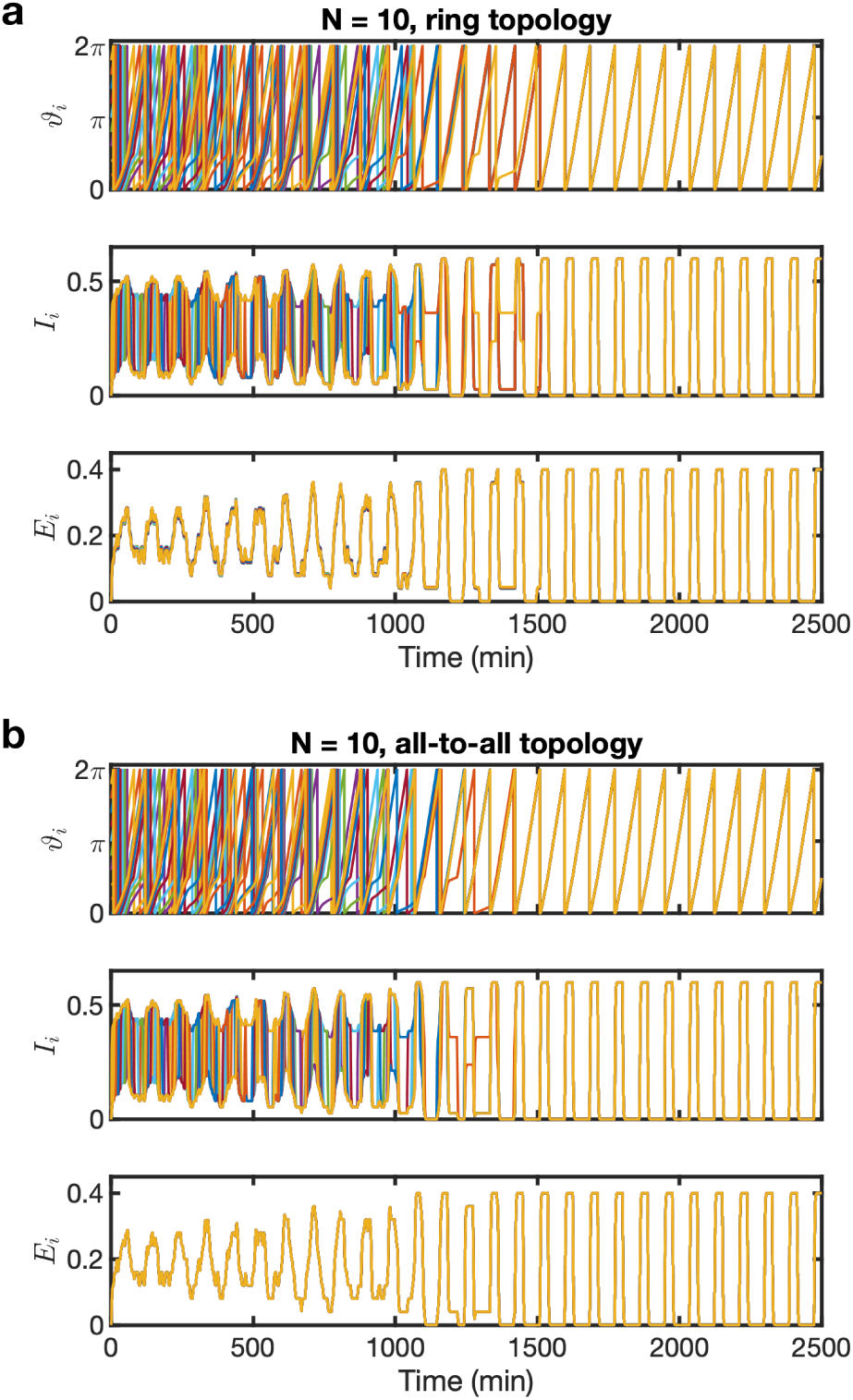
Illustrative numerical simulations considering a population of *N* = 10 cells with both ring (**a**) and all-to-all (**b**) topology. Each numerical simulation depicts temporal evolution of cell-cycle phases *ϑ*_*i*_ (top), internal inducer molecules *I*_*i*_ (middle), and external inducer molecules *E*_*i*_ (bottom).

We then performed a series of more biologically realistic simulations considering a population of *N* = 100 cells. Since the ring topology is not biologically reasonable in this scenario, we validated only the multi-compartment model with an all-to-all coupling. Numerical simulations confirm the results obtained in the scenario with few cells (i.e. *N* = 10). An illustrative numerical simulations is shown in Fig. 4.

**Fig. 4.**
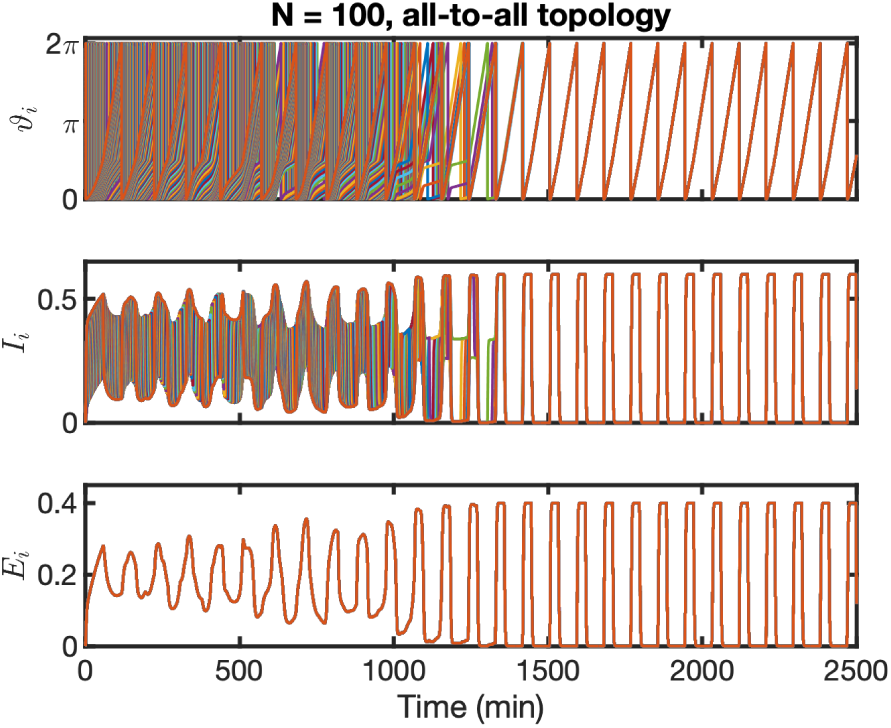
Illustrative numerical simulation considering a population of *N* = 100 cells coupled with an all-to-all topology. The numerical simulation depicts temporal evolution of cell-cycle phases *ϑ*_*i*_ (top), internal inducer molecules *I*_*i*_ (middle), and external inducer molecules *E*_*i*_ (bottom).

### 4.2 Sensitivity analysis

A sensitivity analysis was carried out to assess the robustness of the considered system in achieving a synchronised collective behaviour also in presence of perturbation of the nominal parameters. For simplicity, we consider an ideal quorum sensing control, thus the Hill coefficient in (6) is assumed to be equal to *h* = +∞, and the Hill function in (6) becomes an ideal step.

To evaluate the performance of the proposed system in synchronising the cell-cycle clocks, we introduce the following synchronisation index:

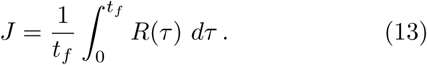

where *t*_*f*_ is the simulation time, and *R*(*t*) is the mean phase coherence of the Kuramoto order parameter (Kuramoto, 1984). The mean phase coherence *R* ∈ [0, 1] is defined as:

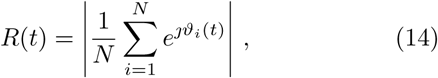

where *ϑ*_*i*_ is the phase of *i*-th cell.

The sensitivity analysis was carried out in the same scenarios used previously to validate the multi-compartment model. The values of the synchronisation index defined in (13) were obtained by varying the external diffusion rate *D* and the dissociation constant *K*. To compute *J*, we consider *t*_*f*_ = 4000 min in (13). The external diffusion rate *D* is varied in the interval [0, 200] min^−1^, whereas the dissociation constant *K* is varied in the interval [0, 0.6[. The choice of the upper bound of the parameter *K* is motivated by the fact that the concentration *I*_*i*_ can not exceed that value, a behaviour that can be appreciated also in the numerical simulations reported in Fig. 3. The results of the sensitivity analysis are shown in Fig. 5a (*N* = 10, ring topology), Fig. 5b (*N* = 10, all-to-all topology), and Fig. 6 (*N* = 100, all-to-all topology).

**Fig. 5.**
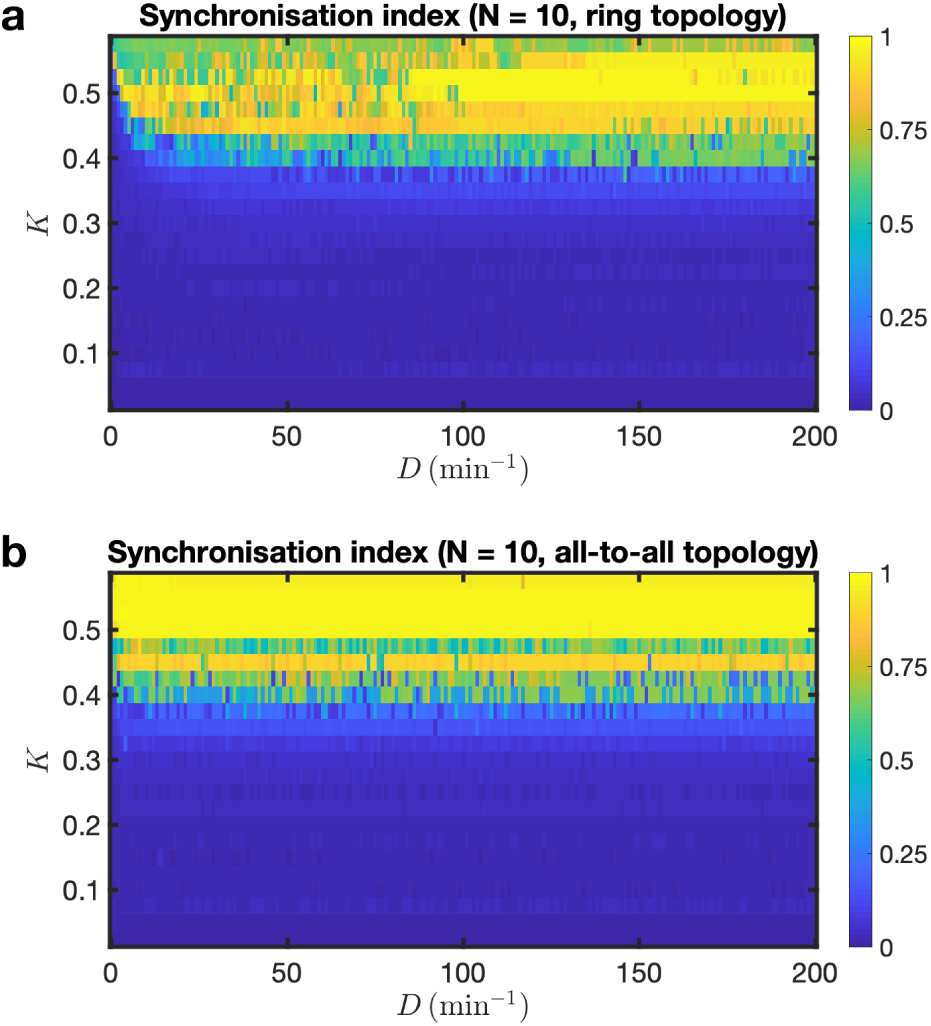
Synchronisation index *J* as a function of the external diffusion term *D* and the dissociation constant *K*, assuming *h* = ∞ in (6). The multi-compartment model was simulated considering a population of *N* = 10 cells coupled with ring (**a**) and all-to-all (**b**) topology.

**Fig. 6.**
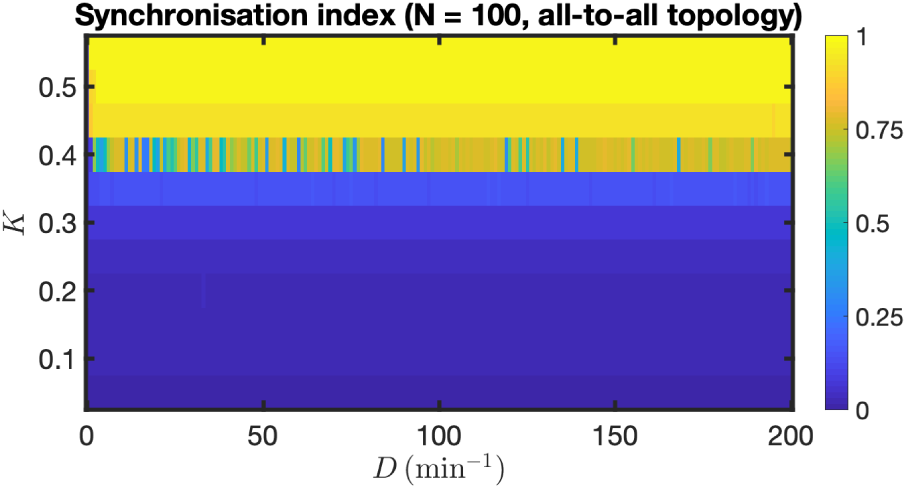
Synchronisation index *J* as a function of the external diffusion term *D* and the dissociation constant *K*, assuming *h* = ∞ in (6). The multi-compartment model was simulated considering a population of *N* = 100 cells coupled with an all-to-all topology.

As depicted in Fig. 5 and Fig. 6, a broad range of perturbed parameters leads to synchronised collective behaviours. In particular, the dissociation constant *K* is the dominant parameter that defines the onset of synchronised collective behaviours. This can be explained by observing that input *u*_*i*_ depends on the concentration of the internal inducer molecule *I*_*i*_. In the assumption of an ideal quorum sensing control (Hill coefficient *h* = +∞), the cell-cycle phase *ϑ*_*i*_ of *i*-th cell is perturbed if *I*_*i*_ is greater than or equal to the dissociation constant *K*. In turn, *I*_*i*_ depends on the concentration of the external inducer molecule *E*_*i*_, that according to the nominal parameters used here has an upper bound equals to 0.4. The latter value is also the minimum value of *K* that leads to synchronised collective behaviours. This means that if the dissociation constant *K* is not large enough, then each cell can produce enough inducer molecule (*I*_*i*_) on its own to start the cell cycle, without having to wait for the external concentration (*E*_*i*_) to increase.

## 5. CONCLUSION

In this work, we have investigated the problem of synchronising the cell-cycle across a population of yeast cells. To address this problem, we proposed the construction of an *in vivo* quorum sensing mechanism to couple the cell-cycle across the cell population. We presented an *in silico* proof-of-principle implementation of the synthetic biomolecular circuit. For the purpose of controlling the cell cycle, we propose to modify the cell-cycle machinery in order to stop the cell cycle in the G1 phase, as done in (Rahi et al., 2016). We also validated the *in silico* implementation through a series of numerical simulations and demonstrated the feasibility of our approach.

To the best of our knowledge, this is the first time that a synthetic quorum sensing system is proposed to couple all the cells in a population to synchronise their cell-cycle clocks. Previously, the synchronisation problem was addressed by considering an open-loop control strategy where an external periodic input was used to entrain the cell-cycle clocks across the cell population, although with limited success (Charvin et al., 2009). Recently, two *in silico* studies have demonstrated the feasibility of using external feedback control of the *CLN2* gene expression to synchronised the cell-cycle clocks (Perrino et al., 2019a,b).

The next step is to implement the proposed quorum sensing system in living yeast cells. A possible *in vivo* implementation can rely on the α-factor mating pheromone, that has been used recently in the design of an orthogonal cell-to-cell communication system in yeast (Shaw et al., 2019).

## ACKNOWLEDGEMENTS

This work was supported by the COSY-BIO (Control Engineering of Biological Systems for Reliable Synthetic Biology Applications) project, which has received funding from the European Union’s Horizon 2020 research and innovation programme under grant agreement 766840.

